# A Model for Navigation in Unknown Environments Based on a Reservoir of Hippocampal Sequences

**DOI:** 10.1101/2019.12.18.880583

**Authors:** Christian Leibold

## Abstract

Hippocampal place cell populations are activated in sequences on multiple time scales during active behavior, resting and sleep states, suggesting that these sequences are the genuine dynamical motifs of the hippocampal circuit. Recently, prewired hippocampal place cell sequences have even been reported to correlate to future behaviors, but so far there is no explanation of what could be the computational benefits of such a mapping between intrinsic dynamical structure and external sensory inputs. Here, I propose a computational model in which a set of predefined internal sequences is used as a dynamical reservoir to construct a spatial map of a large unknown maze based on only a small number of salient landmarks. The model is based on a new variant of temporal difference learning and implements a simultaneous localization and mapping algorithm. As a result sequences during intermittent replay periods can be decoded as spatial trajectories and improve navigation performance, which supports the functional interpretation of replay to consolidate memories of motor actions.

## Introduction

The position of a rodent in a known environment can be decoded from the population activity of hippocampal place cells, which are active at only a few locations of the environment and silent otherwise (O’Keefe and Dostrovsky 1971; Skaggs et al. 1993; Davidson et al. 2009). Much work has been dedicated to successfully unravel neural circuit mechanisms underlying this representation of space (e.g. Touretzky and Redish 1996; Samsonovich and McNaughton 1997; Cutsuridis and Hasselmo 2012; Saravanan et al. 2015; Erdem et al. 2015; Stachenfeld et al. 2017), however, there is still ongoing debate particularly on the mechanisms (e.g. Foster 2017; Liu et al. 2018b; Matheus Gauy et al. 2018; Nicola and Clopath 2019) and the functional role of temporal activity patterns(Liu et al. 2018a). In contrast to technical systems, where autonomous localization methods are based on a costly collection of extensive series of sensory snapshots (Davison et al. 2007; Milford and Wyeth 2012; Siam and Zhang 2017), the prevalent idea in neuroscience is that, in mammals, the hippocampal space code arises from efficient internal dynamics that is locked to places of salient sensory features (Keinath et al. 2018) thereby saving the synaptic resources for explicitly memorizing all details of place-specific sensory inputs. Path integration of the animal’s velocity vector is a widely accepted implementation thereof (Fuhs and Touretzky 2006; Welday et al. 2011; Burgess and O’Keefe 2011; Navratilova et al. 2012) and similar ideas have been proposed for path integration in insects, too (Srinivasan 2015). However, a velocity-based approach precludes mapping of more abstract spaces that are not associated with the physical movement of the animal (Agster et al. 2002; Aronov et al. 2017). An alternative idea to utilize internal dynamics for mapping topological spaces is motivated by extensive reports of hippocampal sequences that are present during locomotion in real and abstract spaces (theta sequences) (Foster and Wilson 2007; Aronov et al. 2017), during wheel running (Nadasdy et al. 1999; Pastalkova et al. 2008; Malvache et al. 2016), and even during offline states (sequence replay) (Lee and Wilson 2002; Davidson et al. 2009; Diba and Buzsaki 2007). Most notably, sequence-like activity patterns were reported to have marked similarity between navigation behavior and offline periods that occur even before the animal navigated a new path (linking of separate experiences) (Gupta et al. 2010), or even a whole new maze for the first time (preplay of future events) (Dragoi and Tonegawa 2011, 2013; Grosmark and Buzsaki 2016; Liu et al. 2018b; Chenani et al. 2019).

There are well-known classical models that offer explanations for place specific firing and partly even remapping on the single cell level, based on either alignment of external reference frames to sensory inputs (Touretzky and Redish 1996; Redish and Touretzky 1997), or combinations of sensory inputs and intrinsic connectivity (Samsonovich and McNaughton 1997). Preplay is not playing a role in these models and, apart from the ongoing debate about the existence of prewired sequences that is held from the perspective of data analytical approaches (Foster 2017), there is the theoretical problem for the preplay hypothesis that there is no account for how these sequences could link to action selection and reward learning in a spatial navigation context. Here, I propose to resolve this shortcoming by introducing a minimal mathematical model of hippocampal sequence propagation, that proposes a theoretical and computational role for hippocampal preplay sequences, in that it connects them to autonomous mapping, and reward-based-learning. The model is based on the assumption that few salient sensory inputs can exert strong hippocampal excitation that drives prewired hippocampal sequences. The resulting theory makes testable experimental predictions for temporal properties of place field activity and the effects of reward on replay.

## Model

A table of parameters (Table 1) is included as reference to the notation used throughout the main paper.

**Table 1:** Summary of mathematical notation.

|  |  |  |  |
| --- | --- | --- | --- |
| $t$ | discrete time | $x$ | state coordinates |
| $N$ | number of possible states | $s_t$ | agent's state at time $t$ |
| $m_i(x)$ | action parameter | $p_i$ | probability for choice of action $i$ |
| $\beta^{-1}$ | temperature of action choice | $b(x)$ | familiarity of state $x$ |
| $F$ | Number of features | $f_i$ | feature state indices |
| $r_j(x)$ | reward for action $j$ in state $x$ | $\varepsilon$ | learning rate |
| $v(x)$ | reward prediction at $x$ | $w_n$ | reward prediction at state $n$ |
| $\delta$ | reward prediction error | $\delta_{nm}$ | Kronecker delta |
| $\ell_n$ | memory trace for $w_n$ | $\gamma$ | decay rate of $\ell_n$ |
| $\mathcal{S}_f^{(i_f)}$ | $i_f$-th copy of $f$-th sequence | $\sigma_f^{(i_f)}$ | state of sequence $\mathcal{S}_f^{(i_f)}$ |
| $L$ | sequence length | $R$ | ensemble repetitions in sequence |
| $\mathcal{X}(t)$ | ensemble activity matrix | $J$ | linear decoder |
| $\hat{s}$ | state estimate | $K$ | state encoder |
| $h$ | sensory input | $p_j^*$ | choice probability including exploration |
| $p_e$ | exploration probability | $\tau_e$ | exploration decay constant |

How a set of prewired hippocampal sequences can be used to aid navigational learning is first illustrated for a schematic maze with an agent whose state, for now, is solely determined by the position in the maze (Figure 1A), i.e., the number of the node *n* = 1, …, *N*. The agent is navigating the maze in discrete time. The Euclidean coordinates of the agent at the integer time step *t* is contained in the vector *s*_*t*_. The agent moves according to a probabilistic choice between at most 4 possible actions *i* = 1, …, 4 (’Go North’, ‘Go West’, ‘Go South’, ‘Go East’) governed by choice probabilities

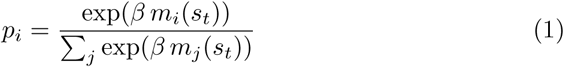

derived from plastic action parameters *m*_*i*_(*x*_*n*_) at node *n* where the 2-dimensional Euclidean coordinate vector is denoted as *x*_*n*_. The action parameters *m*_*i*_(*x*_*n*_) are changed according to a variant of reinforcement learning introduced in the subsection “Reinforcement Learning” below. Simulations were carried out at the inverse temperature *β* = 5.

**Figure 1:**
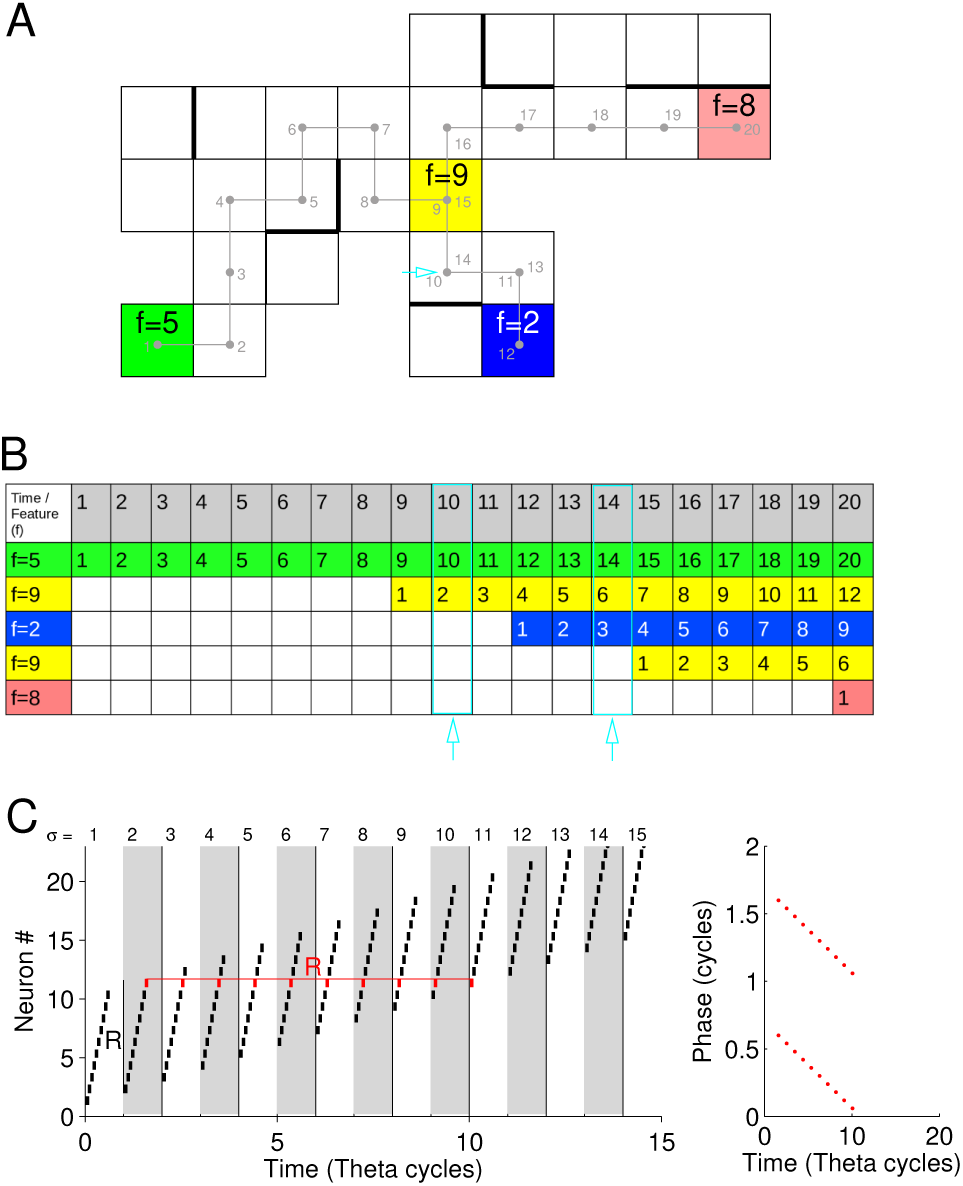
Model outline. (A) An agent navigates a 2-d maze (trajectory and time steps in gray; thick lines indicate walls). Few of the nodes are selected as salient locations (colored squares, only four shown *f* = 2, 5, 8, 9) (B) Time evolution of sequences (grey row indicates discrete time). At feature nodes a sequence starts whenever the agent visits the node (numbers indicate sequence state). The position of the agent can be decoded by the state vector (column) in the sequence table. Single nodes can be represented by multiple state vectors (e.g. cyan arrows). (C) Interpretation in terms of neuronal ensembles. Each sequence state *σ* is thought to reflect neuronal activity in a theta cycle (grey and white rectangles). Neuronal spikes (black ticks) are representative for active ensembles. Each sequence state *σ* is realized by a short theta sequence of *R* ensembles. Every ensemble is thus active *R* times (red horizontal line). Sequential activity generates phase precession (left: red dots represent phases of red spikes on the left; every spike is plotted twice).

Two technical measures were implemented to facilitate exploratory behavior (and reduce simulation time). First, a familiarity term *b*(*x*_*n*_) is subtracted from all action parameters *m*_*i*_ that connect to node *n*. The familiarity *b*(*x*_*n*_) is increased by +1 every time the agent’s internal estimate *ŝ* (see below) of its current position equals *x*_*n*_. After that all familiarities *b*(*x*_*n*_) are reduced in every time step by a multiplicative factor *e*^−1*/*50^ ≈ 0.98, unless otherwise mentioned. Mechanistically, familiarity implements a memory about the novelty of local sensory inputs (Brady et al. 2008) the importance of which for navigational decisions decays with a time constant of 50 time steps. Most importantly, familiarity does not give information about the relative spatial relation between different nodes, and thus does not allow to construct a spatial map. The inclusion of the familiarity bias is thus solely to impose a repulsive force to avoid revisiting recently seen locations.

Second, to avoid agents cycling between two nodes the movement is restricted in that a penalty term *c* = −10 is added to the action parameter *m*_*i*_ that leads back to the node *s*_*t*−1_ the agent has visited in the previous time step. This penalty term is then deleted in time step *t* + 1.

### Feature nodes

Initially, simulations are conducted for symmetrically connected 2-dimensional random graphs generated as described in the Appendix E. Within a maze *F* « *N* feature nodes *f*_1_, …, *f*_*F*_ ∈ {1, …, *N*} are randomly selected such that 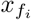 and 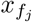 are separated by a Euclidean distance of at least 3 for *i* ≠ *j* (for simulations with *F/N* ≥ 0.2 the minimal distance between feature nodes was set to 2). In the first set of simulations, mazes are considered to have sparse feature density in which only in *F/N* = 5% of the nodes a sensory cue (feature) is placed that is salient enough to evoke activity in the hippocampus starting one of the prewired sequences (Figure 1B). The selection of which specific sequence is evoked by a feature is assumed to be random and may, e.g., be explained by fluctuations of cellular excitability. Sequences thereby get dynamically attached to sensory experiences. The agent is assumed to have correct memory about the identity of the feature nodes and is able to identify them when it is located at one of those without error. In this way sequence activation at the feature nodes (via extrahippocampal pathways) implements the alignment of sensory signals with extrahippocampal coordinate systems as supposed in previous non-sequencebased models (Redish 1999).

In each maze, one randomly chosen feature node is selected as goal location (yellow triangle in Figure 1). A reward *r*_*j*_(*s*_*t*_) = 10 is delivered to the agent once it reaches the goal location at time step *t* + 1 from node *s*_*t*_ by taking action *j*.

### Sequences

While navigating the maze, the agent propagates internal activity sequences, reminiscent to hippocampal theta sequences (Foster and Wilson 2007; Feng et al. 2015). There are as many different sequences as there are feature nodes and the sequence 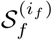 associated with feature *f* = 1, …, *F* is activated and set to state 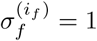 whenever the agent visits the feature node *x*_*f*_. The upper index *i*_*f*_ counts the active instances of the sequence *f* and is increased by 1, whenever the agent visits the node *x*_*f*_ again. In each time step all active sequences are propagated by 1, i.e.

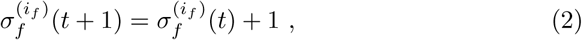

implying that sequence propagation scales linearly with running speed as suggested by speed-controlled hippocampal phase precession (Geisler et al. 2007). Whenever an instantiation of a sequence reaches state *σ*_*f*_ = *L* (the sequence length) it is inactivated. Note that, implementing sequence propagation in this way, a sequence can be active multiple times (indicated by the instance counter *i*_*f*_) in different states and, consequently, the same position can be represented by multiple states of the sequence (cyan arrows in Figure 1B).

Biologically, sequences correspond to the successive activation of neuronal ensembles during the theta state (Foster and Wilson 2007; Feng et al. 2015). Each ensemble can be active during multiple successive theta cycles. Each state *σ* is therefore identified by a set of *R* ensembles played out in sequence during a theta cycle; Figure 1C. The number *R* then also corresponds to the number of theta cycles an ensemble is repeated, i.e., the ensemble starting the sequence of *R* patterns in sequence state *σ* is active min(*R, σ*) − 1 times before the sequence is in state *σ*; red spikes in Figure 1C. A suggestion for a potential neurobiological implementation of such a link between movement and theta sequences is outlined in Appendix A.

Formally, the sequences that define the internal state of the agent are translated to a binary matrix 𝒳 (*t*) ∈ {0, 1}^*F* ×(*L*+*R*−1)^ indicating all active ensembles at time *t*:

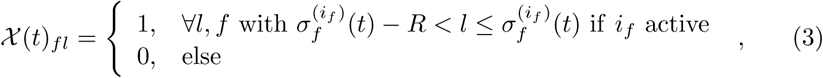

with *f* identifying the feature node and *l* the ensemble starting the sequence at state 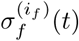. The internal dynamics captured by the state matrix 𝒳 is illustrated in Figure 1B.

### Encoding and Decoding Space

The state matrix 𝒳 links the sensory experience of the agent at the feature nodes with the internal dynamics and thus it allows to make positional inferences also for nodes *x*_*n*_ without features. To do so, a decoder *J* ∈ ℝ^*N*×*F* ×(*L*+*R*−1)^ is trained such that it maps the internal state 𝒳 (*t*) to a positional estimate *ŝ*(*t*) by

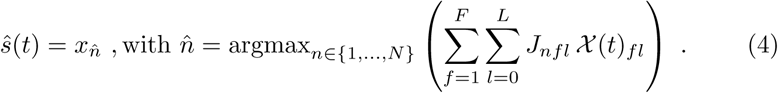

The decoder is trained using a simple conditional increment

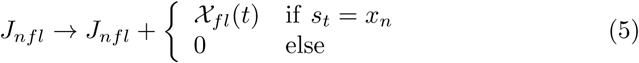

i.e., the readout weight from sequence state (*f, l*) to position *n* is increased if and only if this sequence state was active at position *n*. After each increment the updated row is normalized to 1:

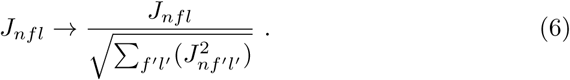

A new row index *n* of the decoding matrix *J* is generated, every time the agents hits a node for the first time. This procedure assumes that all nodes are sufficiently dissimilar to identify novelty. In principle the model could also implement confusions, i.e., same decoder rows refer to multiple physical maze locations, which is not explored in this paper.

### Reinforcement Learning

Navigational learning on the maze is implemented by changing two node-specific entities, the reward prediction weight *w*_*n*_ and the action parameters *m*_*j*_(*x*_*n*_) of node *n* by reinforcement learning, whereas the sequence generating network is assumed to remain constant.

In contrast to standard reinforcement learning (cf. Dayan and Abbott 2001) of navigational strategies, the maze is initially unknown to the agent, i.e., the positions *x*_*n*_ do not yet have an internal representation but need to be estimated 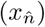 from previous experience by decoding from the states of all active sequences (see previous subsection). The learning therefore takes place at the weight 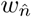 and action parameters 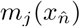 of the estimated location. The changes of *w*_*n*_ and 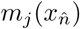 are driven by rewards *r*_*j*_(*s*_*t*_) delivered to the agent for performing action *j* at the actual position *s*_*t*_. More specifically, if at time step *t* the agent decided to perform motor action *j*, the action variable is updated according to

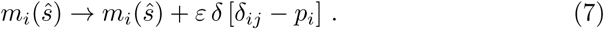

After each update, the action variables at an estimated node are normalized to length 1. In equation (7), *δ*_*ij*_ is the Kronecker-symbol, *p*_*i*_ is the action probability obtained via equation (1), *ε >* 0 is a fixed learning rate parameter, and

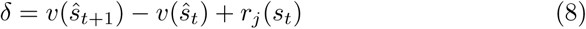

is the reward prediction error (Sutton and Barto 1990; Suri and Schultz 1998), not to be confused with the Kronecker delta (Dayan and Abbott 2001). The reward prediction error *δ* consists of the actual reward *r*_*j*_(*s*_*t*_) and the difference *v*(*ŝ*_*t*+1_) − *v*(*ŝ*_*t*_) between the agent’s reward predictions based on the internal positional estimates. The reward predictor

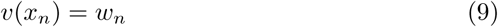

is learned such that, in each time step, the weights *w*_*n*_ are updated using a *synaptic memory trace 𝓁*_*n*_ according to

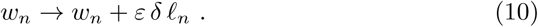

Thereby, the memory *𝓁*_*n*_ is incremented by 1 whenever the internal positional estimate 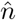 equals *n* and otherwise decays with rate 0 *< γ <* 1:

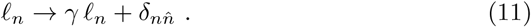

The implementation of the weight update via a synaptic memory trace is nonstandard in reinforcement learning, but it serves a purpose similar to the eligibility trace in temporal difference learning (Sutton 1988; Dayan 1993) where an increased spatial spread of the reward information is usually obtained by low-pass filtering the state sequence *s*_*t*_. In the present model there is only an estimate of the state available and, also, internal activity was reserved exclusively for sequence propagation. I thus decided to implement the eligibility trace within the synaptic plasticity rule, as know from models of synaptic meta-plasticity (Fusi et al. 2005; Ben Dayan Rubin and Fusi 2007; Leibold and Kempter 2008; Päpper et al. 2011).

All simulations in this paper have been performed for decay of *γ* = 0.75. The effects of changing *γ* on learning performance are discussed in the Appendix B, in which the most important result is that replay (see below) allows for much shorter decay constants to achieve similarly high performance. All simulations were performed with a learning rate *ε* = 0.025.

### Replay

After the agent has reached the reward location (end of a behavioral trial), a replay epoch is started, except for simulations without replay. Unless mentioned otherwise, during each replay epoch the model iterates through all feature nodes the agent has visited in the previous exploratory trial and starts the sequences attached to the respective feature nodes one at a time (as in contrast to the behavioral trial where multiple sequences are active simultaneously). In each time step the linear decoder *J* is used to estimate a position *ŝ*′(*t*) from the internal state matrix 𝒳′ (*t*) that propagates only the one sequence for the current feature node. Once the sequence has reached its end state *σ*_*f*_ = *L*, a new sequence is started that belongs to the feature node visited next during exploration.

While the sequences are propagated, in each time step reward prediction errors 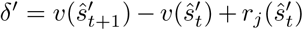 are computed and used to update the action parameters *m*_*i*_(*x*_*n*_) but not the weights *w*_*n*_.

The update of the action parameters follows the classical learning rule

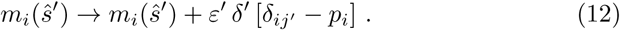

where the replayed action *j*′ is determined by the largest overlap between the estimated movement vector 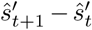 and the direction implicated by the action in 2-dimensional Euclidean space. The learning rate during replay is increased to *ε* = 0.125.

### Open Field

In contrast to the previous simulations the state variable of the agent now also includes heading direction *h*. Eight heading directions are considered (North, Northeast, East, Southeast, South, Southwest, West, Northwest). As a consequence of including heading the motor actions are reduced to “Move forward”, “Turn left by 45 degrees”, and “Turn right by 45 degrees”. Including heading also allows to define direction dependent features as they would be induced by e.g. distant landmarks. The state dynamics are applied to position and heading in the same way as for the four possible movements in the 2-d random mazes.

An quadratic environment of *N* = 11 × 11 nodes was set up including 37 feature nodes. Five of the feature nodes (corners and center) were defined direction invariant (sequence is started independent of *h*). At twelve feature nodes (three per corner at *L*^∞^-norm distance 1) the sequence is only triggered for outbound headings. At another twenty features (five per corner at *L*^∞^-norm distance 2) sequences are triggered for all inbound headings.

#### Firing Rate Maps

Since no individual neuron dynamics are simulated but only sequences of pooled cell ensembles, place fields can only be derived on a single ensemble basis. Each ensemble is uniquely identified by the combination (*f, σ*_*f*_) of sequence number *f* = 1, …, *F* and the state *σ*_*f*_ = 0, …, *L*.

To identify place fields, first firing rate maps were computed for each ensemble by counting how often it was active at a particular node during navigation and this count was divided by the occupancy of the node, i.e., how often the agent visited the node. Nodes that were visited less than 5 times during all trials were excluded from the analysis.

To compute place field statistics for open field simulations, standard experimental procedures were followed (Leutgeb et al. 2004) and firing rate and occupancy maps were additionally smoothed by a Gaussian kernel with standard deviation 1 (spatial bin).

#### Direction tuning

To assess directional tuning of place fields, a maximum likelihood model (MLM) algorithm was used that has been first introduced in (Cacucci et al. 2004). In short the algorithm iteratively fits the place field activity with a multiplicative model in which the firing probability is modeled by a product of a positional term and a directional term. The directional term from the maximum likelihood solution is used for directionality plots (polar plots in Figure 5A) and for computing Rayleigh vectors.

**Figure 2:**
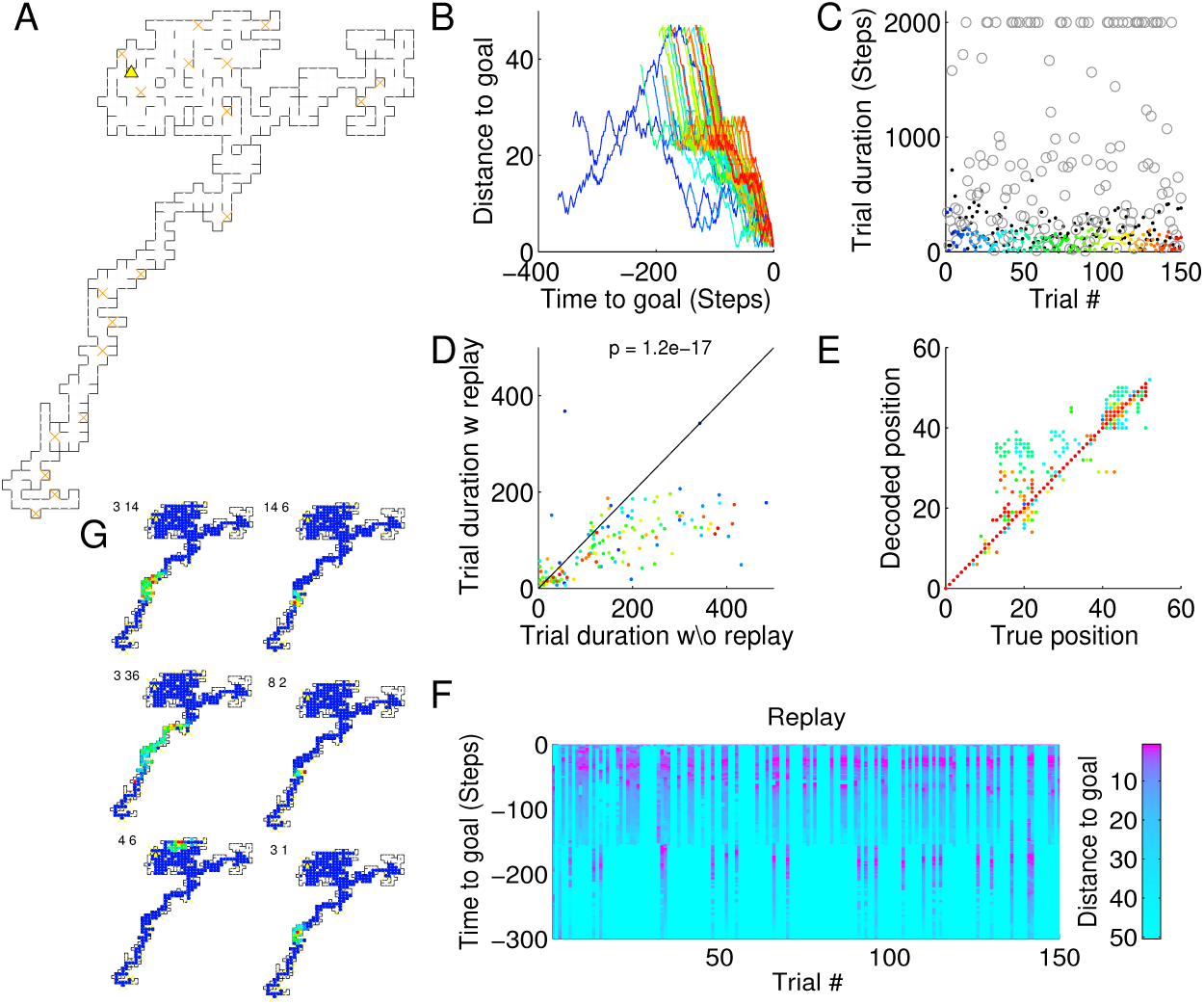
Example session. (A) Example random maze used in the simulations for panels B-G. Brown crosses are feature nodes, yellow triangle marks goal location. (B) Agent learns to approach the goal location (blue: early trials, red late trials). Note that start locations are random and thus lines start a different distances; parameters were *R* = 7, *L* = 150. (C) Trial durations (trials are truncated after 5 × *N* = 2000 time steps). Colors as in B. Grey circles are from random walks starting at the same initial condition, black dots are from simulations without replay. (D) Replay improves navigation (signed rank test). Colors as in B. (E) Decoding of positional coordinates (2 per node) improves with trials. Colors as in B. (F) Replay sequences can be spatially decoded. (G) Examples of place fields. Numbers at the bottom left of each plot are ensemble indices (*f, l*) as in equation (3).

**Figure 3:**
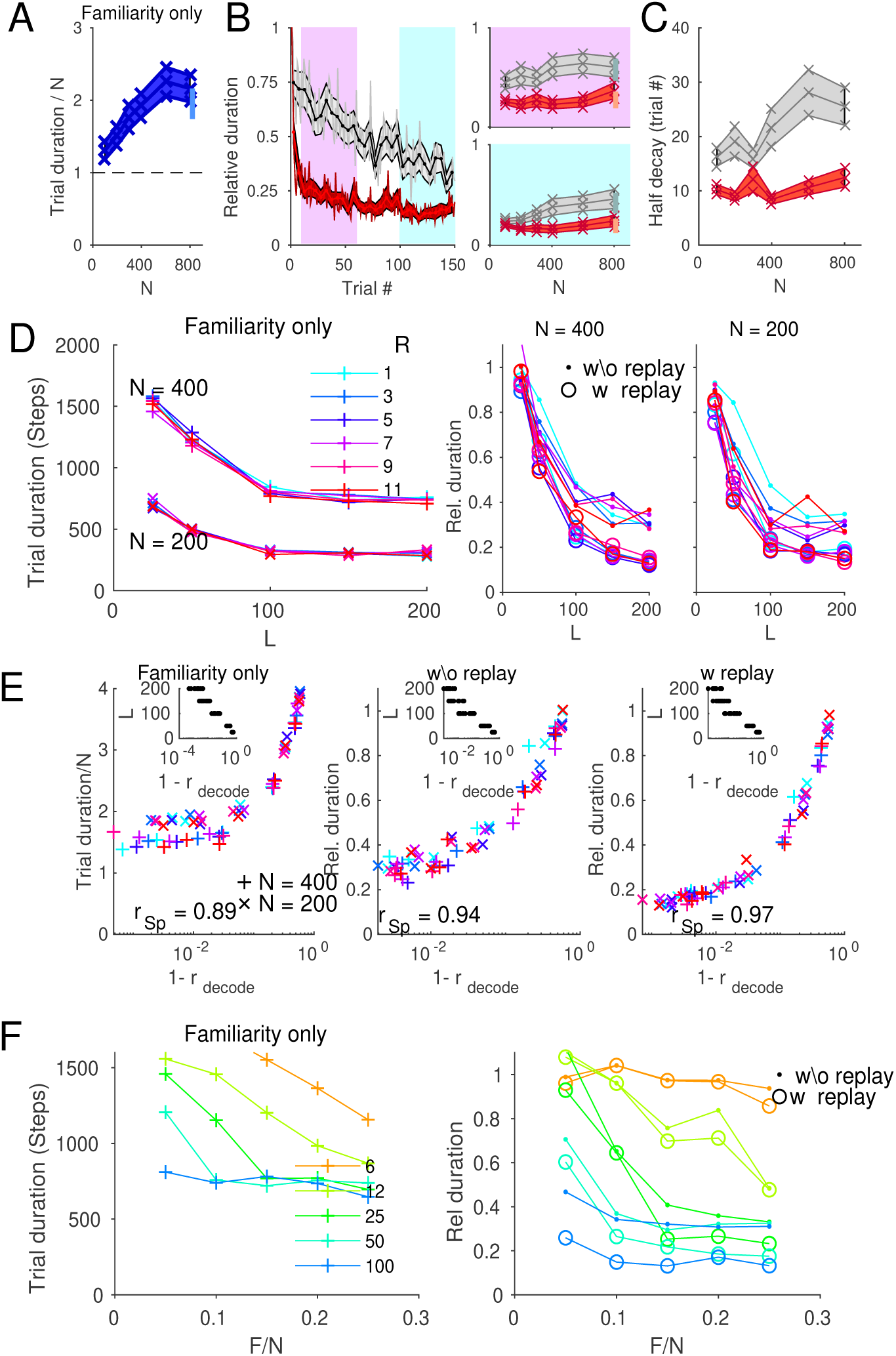
Learning curves and performance measures. (A) Mean ± s.e.m. of trial durations as a function of the maze size *N* for 50 random mazes without reinforcement learning (familiarity only). The light blue bar at *N* = 800 indicates mean ± s.e.m for a slower familiarity memory time scale of 100 time steps. (B) Left: Relative duration (for every maze, duration with reinforcement learning is divided by duration without reinforcement learning) as a function of trial number. Grey traces indicate mean ± s.e.m for simulations without replay epochs, Red traces are derived from simulations with replay epoch after each trial. Pink box marks trials 11 to 60, cyan box marks trials 101 to 150. Right: Mean ± s.e.m. averaged over cyan (bottom) and pink (top) interval as a function of maze size *N*. Light symbols at *N* = 800 stem from simulations with familiarity memory time scale of 100 time steps. (C) Decay rates (slope of relative duration at 10 trials) for simulations with (red) and without (grey) replay. (D) Relative trial durations as functions of sequence length *L* and repetition number *R* (color code). Left: mean durations without reinforcement learning for two maze sizes *N* = 400 and *N* = 200. Right: relative durations for simulations with replay (circles) and without replay (dots). Graphs in left and right panel are obtained for a different mazes size *N* as indicated. (E) Durations as a function of decoding error (1 − *r*_decode_, where *r*_decode_ denotes the Spearman correlation coefficient between true and estimated position coordinates, *s* and *ŝ*, respectively). The rank order correlations *r*_Sp_ indicate inverse correlations between decoding error and search performance. Insets: sequence length *L* and decoding error are negatively correlated. (F) Durations as a function of feature density *F/N* for different sequence lengths *L* (indicated by different colors). Crosses: no reinforcement learning; dots: reinforcement learning without replay; circles: reinforcement learning with replay.

**Figure 4:**
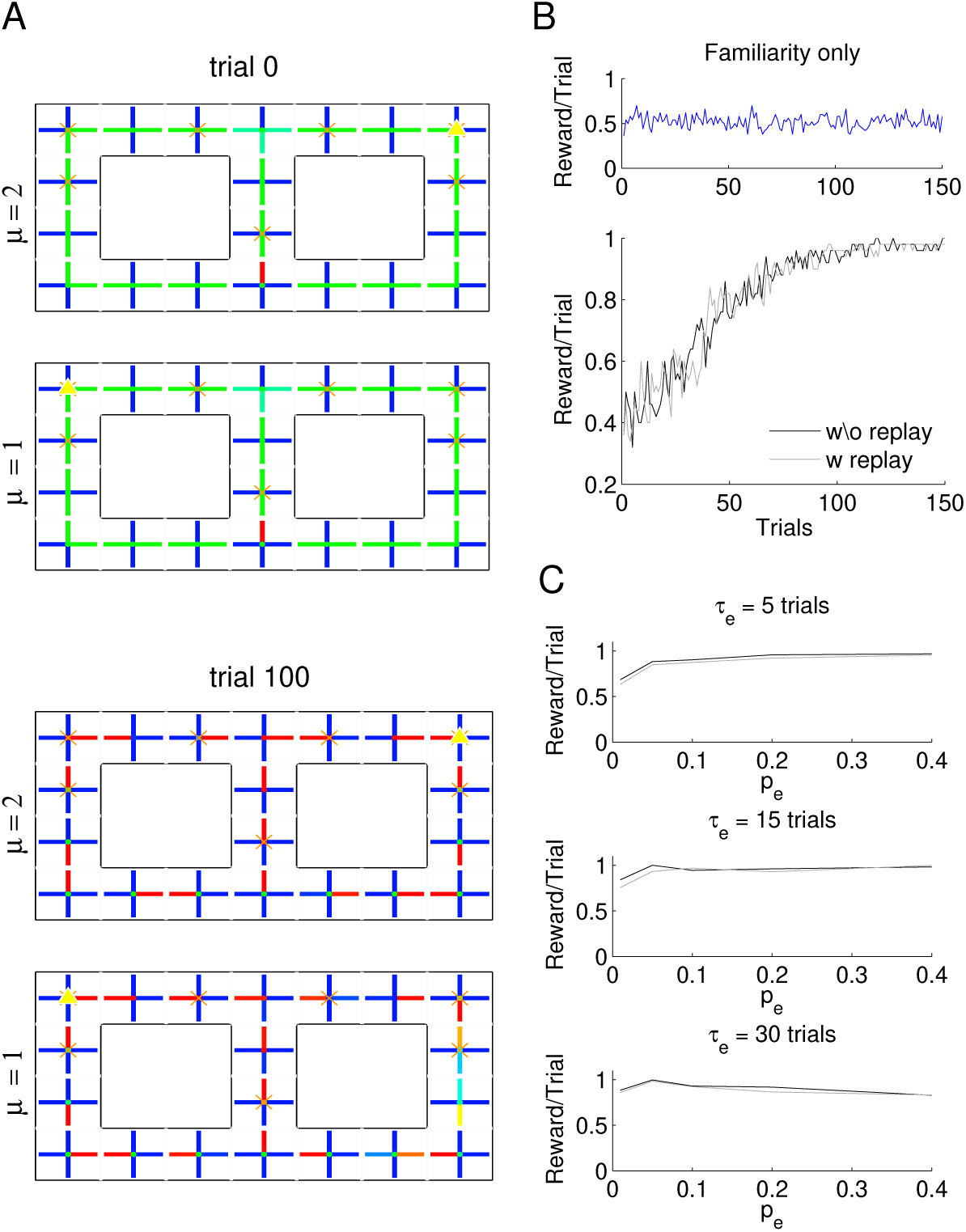
Internal states. (A) Figure-8-maze with initial configuration (top) and final configuration (bottom). Feature nodes are marked with yellow triangles. Each trial is started in the bottom node of the central arm, and the agent leaving this node upwards. Each maze is represented as many times as there are internal motivational states *µ* = 1, 2 (rewards indicated by triangles). In the example shown, the first state is learned to be associated with a reward in the left upper corner, the state *µ* = 2 is learned to be associated with a reward in the right upper corner. Color of ticks indicates the strengths of action parameters *m*_*j*_, the colored dots in the node centers show the learned reward expectation weights (red: high, blue: low). (B) Learning curves (mean of 50 repetitions) for simulations without reinforcement learning (top: 0.5 is chance) and with reinforcement leaning (bottom: results from simulations with and without replay are depicted by different gray levels). (C) Performance depends on exploration parameters (see text). Grey levels as in B.

**Figure 5:**
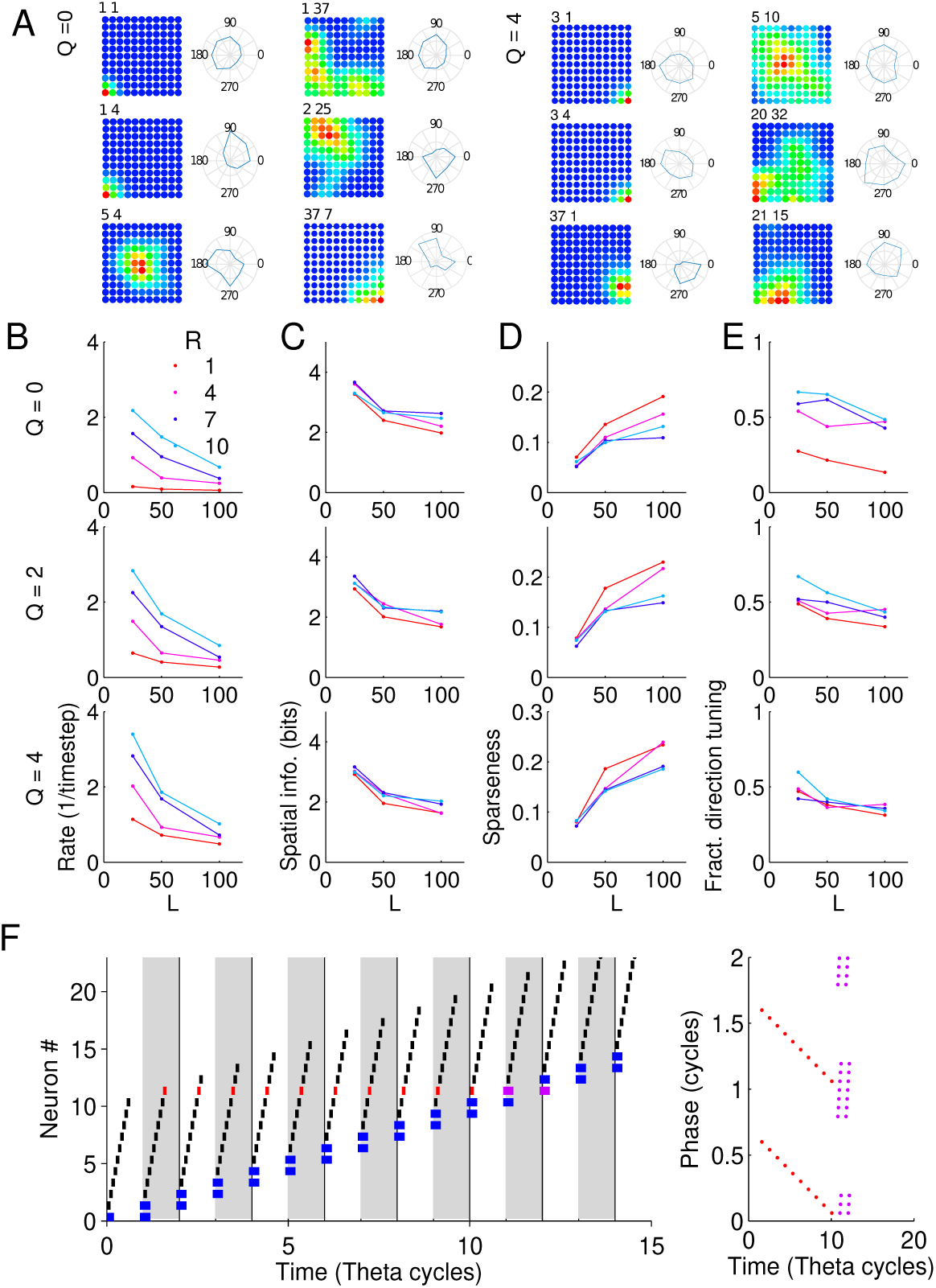
Place fields and direction tuning. (A) Examples of place fields and direction tuning (polar plots) for simulations without sensory spikes (*Q* = 0, left) and simulations with sensory spikes in four subsequent cycles (*Q* = 4, right). (B) Mean peak firing rates as a function of sequence length *L*, repetition number *R* (colors) and number of sensory cycles *Q* (rows). (C) Spatial information (arranged as in B). (D) Sparseness (arranged as in B). (E) Fraction of significantly direction tuned fields (arranged as in B). (F) Schematic illustration of sequences; see Figure 1C. Black ticks indicate spikes during prospective sequences. Blue bars indicate high probability for sensory-evoked activity. The number of black spikes in a theta cycle (gray and white rectangles) equals the number *R* (here 11) of how many theta cycles a cell is active during a sequence. After a cell has fired *R* times it can be activated *Q* (here *Q* = 2) additional times by sensory inputs which is to weak or imprecise to trigger the full sequence. The red (purple) spikes are translated into a phase precession plot (right) for the specific neuron # 12. The purple dots mark the range of phases where sensory spikes might occur.

Place fields are called directional if their Rayleigh vector length (RVL) exceeds the 95-percentile of RVLs from control simulations simulations in which the maze was randomly rotated every 100 time steps and heading was evaluated according to the original maze orientation.

#### Sensory learning and sensory activation

In open field simulations the impact of additional sensory-evoked activity on place fields at non-feature nodes was evaluated. Such activity was generated when the sensory input to ensemble (*f, l*)

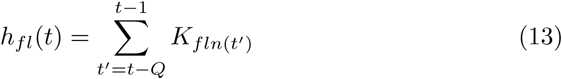

exceeded a threshold of 0.5 (*f* is the feature index, *l* is the ensemble index); Figure 5F. The repetition number *Q* counts how many time steps after the agent has visited location *x*_*n*_ the sensory input is active.

The matrix *K* was learned during navigation according to

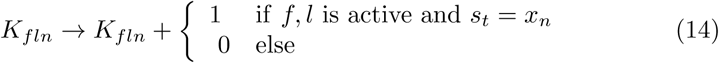

After every update the rows of *K* are normalized to 1. The matrix *K* thus implement a type of “pseudo inverse” to the decoding matrix *J*.

### Simulations

Navigational learning for an example search task (maze with *N* = 400 nodes shown in Figure 2A) in which the agent has to approach a fixed reward location (triangle in Figure 2A) from a random start location in a maze is summarized in Figure 2B, C (see also Supplementary Movies 1&2 and Appendix F for description). While in early trials it takes longer until the agent finds the reward, late trials end successfully in typically less than 150 time steps. Since starting locations are random the trial durations are compared to a random search with only familiarity memory but no reinforcement learning (control condition gray circles) and a version of the model without replay (black dots). Replay improves the navigation performance significantly (signed rank test, *p* = 1.2 · 10^−17^, trials are initialized at identical start positions) by about a factor of two (Figure 2D). Particularly in late trials the replay version in this simulation always outper-formed the non-replay version. The improvement of navigational performance is thereby associated with an improvement of the agent’s ability to estimate its location during the search (Figure 2E); a more detailed analysis of decoding errors will follow later (Figure 3E). Hence, while (theta) sequence activity during search implements a neural representation of the agent’s spatial position as a basis for decision making, offline replay seems to facilitate action learning similarly to memory consolidation.

Sequence propagation during replay could be decoded in terms of spatial trajectories (using the learned linear decoder *J*) which often appeared as a spatially continuous approach to the goal location (smooth color transitions in Figure 2 F; see Supplementary Movies 3& 4) and could thus be interpreted as fast replay of spatial trajectories, although the sequences were wired up already before the maze learning started.

Since sequences were elicited by spatially locked features, neuronal ensembles in these sequences were preferentially activated in the vicinity of their feature locations. The standard procedure that is used to compute place maps (see Methods and, e.g. (Leutgeb et al. 2004)) was then applied to the ensemble activations and the resulting firing maps had qualitative similarities to hippocampal place fields (Figure 2G). Place field properties are going to be investigated in greater detail for open field simulations below.

### Parameter dependencies

So far, all results have been obtained for one specific realization of the maze. Next, it is shown that the results generalize to different mazes by repeating all simulations for 50 realizations of the maze-generating process in each parameter regime. For each maze the simulation without reinforcement learning, just relying on familiarity memory (Figure 3A) is taken as the control condition. For a fixed memory time scale of the familiarity trace, the trial duration for the familiarity-only search strategy was up to about 2.5 times larger than the maze size and thus highly inefficient. However, increasing the memory time scale for familiarity in larger mazes can moderately but non-substantially reduce the trial duration.

The trial durations for model variants with reinforcement learning were then compared to the corresponding familiarity-only simulations (same maze, same initial condition, same random seed) and the relative duration of the trial (duration with learning divided by duration without learning) was used to quantify navigational learning performance (Figure 3B, left). In the case without replay (and *N* = 400), trial duration decreases with experience and relative duration approaches about 0.4 after 150 trials for the used parameter set. In simulations with replay, relative duration values not only decreased to about 0.2 after 150 trials, but most importantly the convergence rate is much higher as evidenced by a comparison of the mean relative duration in trials 10 to 60 (Figure 3B right, pink, Wilcoxon rank-sum test: all p values below 0.0002) and in trials 100 to 150 (Figure 3B right, cyan, Wilcoxon rank-sum test: all p values below 0.0013). The relative duration averaged over early and late trials are relatively independent of maze size *N*. Only for small mazes (low *N*) there is little effect of replay on the asymptotic performance (Figure 3B right, cyan). Thus the benefits of replay seem to be particularly strong for large and complex problems. The speed of convergence, however, is profiting from replay for all investigated parameter regimes (Figure 3C). Number of trials until relative duration crossed a value of 0.5 was significantly smaller with replay for all maze sizes (Wilcoxon rank-sum test: all p values were below the value of 0.0016 obtained for the smallest maze *N* = 100).

To understand what are the essential mechanisms that allow the model to work, the learning performance was monitored for the two parameters sequence length *L* and the number *R* of repetitions of an ensemble in subsequent time steps (Figure 3D,E). Whereas the repetition number *R* does not have a systematic and large effect on the trial duration in neither the familiarity-only control, nor in the reinforcement learning models, all performance parameters crucially improve with *L*, leveling out at an asymptotic value of *L* ≈ 100 (Figure 3D). The sequence length *L* also negatively correlates with the decoding error 1 − *r*_decode_ (insets in Figure 3E) and thus the decoding performance was the fundamental predictor for the success of navigational learning (Figure 3E).

So far, features have been distributed very sparsely in the simulated mazes, and thus sequences were required to bridge long distances from one feature node to the next. It could therefore be suspected that the long sequence length *L* ≈ 100 needed to reach asymptotic performance reflects this sparse ornamentation of the maze. To directly test this assertion, navigational learning was investigated in mazes with higher feature loads *F/N* up to 25% (Figure 3F).

As expected, an increase in the feature load *F/N* improves the performance parameters for short sequence lengths *L* but not the asymptotic values at *L* = 100. The numerical results suggest that sequences of length *L* ≈ 25 may still be sufficient for good navigational learning if the maze has sufficiently many feature nodes, whereas long sequences need to be used in maze with only few landmarks.

While the random mazes used in the previous paragraphs have served well to investigate the basic parameter dependencies of the model, they miss out important aspects of a navigation task for real animals, such as heading direction, distal cues, and internal motivation. For the next set of simulations, I therefore extended the model design to be able to also capture these aspects and thereby focused on two classic behavioral paradigms used in rodent research, the figure-8 alternation task (Wood et al. 2000; Ainge et al. 2007) and a free foraging task in an open arena (Muller et al. 1987; O’Keefe and Burgess 1996).

### Figure-8-Maze

In the figure-8 alternation task an animal has to alternate between two arms following traversal of the stem section that needs to be passed in every trial (Figure 4A). Animals get rewarded on their way back to the start location only if they made the correct choice and then continue to run to the start location for the next trial.

To model this task the maze structure has been altered as compared to the random mazes in three ways. First, features are not placed at random but at 7 locations that are salient in the behavioral context (Figure 4A). Second, the state space as well as the action space have been extended by an internal motivation state variable *µ* to distinguish between the intentions of a leftward and a rightward run. More specifically, the action parameters *m*_*i*_(*ŝ, µ*) and reward prediction *v*(*ŝ, µ*) are updated for each internal state *µ* independently. In contrast to the position that, again is estimated from a learned decoder, the motivational state is assumed to be exactly known to the agent. Furthermore, the model was extended by an exponentially decaying initial random exploration phase to force the agent to search for a second reward even if one reward position is already known. Exploration is defined by two parameters, the initial exploration probability *p*_e_ and the decay time constant *τ*_e_. The probability of an action *j* was modeled as 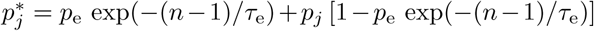, where *p*_*j*_ is the original probability derived from the action parameters *m*_*j*_(*x,µ*) from equation (1), and *n* is the trial number.

Besides the four possible actions related to spatial movement the model now also includes an action for each transition between internal states. The number of motivational states is thereby considered to be dynamical and equally so the number of possible actions related to state transitions. The simulations start with no motivational state present. Whenever the agent hits an unknown reward location for the first time, a new motivational state is generated including the corresponding action parameters *m*_*i*_ that govern the transitions to and from all previous states to the new state. In this way the agent develops motivation-dependent spatial activity known as splitter cells (Wood et al. 2000) without explicitly including pre-knowledge of the task.

After the agent has received a reward from a known state the motivational state is forced to change to another state using the momentary values of the action parameters *m*_*i*_ for state transitions (excluding the transition to the current state).

Figure 4B depicts the mean learning curve for 50 repetitions of the task with different random seeds and shows that the agent learns to perfectly alternate after about 100 trials (an example session is shown in Supplementary Movie 5). Performance is independent of replay arguing that path integration is not the major problem but rather the switching between motivational states. The initial exploration phase turned out to be crucial for achieving close-to-perfect performance (Figure 4C) but the dependence on its parameters is rather weak. For decay time constants *τ*_*e*_ of 10 to 30 trials the agent learns the task perfectly for a wide range of initial exploration probabilities *p*_*e*_ ≥ 0.05.

### Free foraging in open fields

For the next set of simulations, I altered the model design to be able to also capture head directionality and thereby focused on a classic behavioral paradigm used in rodent research, the free foraging task in an open arena (Muller et al. 1987; O’Keefe and Burgess 1996). To this end, the model was amended to more closely fit the spatial extent of the animal by introducing eight heading directions *h* (spaced 45 degrees) and by allowing only three motor actions (forward, left-turn, right-turn). The free foraging task does not imply specific reward locations, the animals are rewarded at random times independent of their location just to keep them moving. For a reinforcement learning model such random reward presentation is equivalent to not learn reward expectation weights at all, since reward expectation is equal at all locations and therefore the reward expectation error *δ* is zero. The simulations are thus just done to generate place fields from sequence activity that is locked to salient locations. To this end, the quadratic open field (*N* = 11 × 11 nodes) is ornamented with 37 symmetrically distributed features that are both place and heading dependent; see Methods Open Field. In brief, five of these features (corners and center) are heading invariant all others have heading dependence (inbound to and outbound from corners). Some features are placed on multiple neighboring nodes (inbound to corners). The choice that inbound and outbound trajectories relative to the corners are salient features is of course highly speculative and thus this set of simulations can only be considered as proof-of-principle asking for future indepth analysis of how the spatial distribution of features influences place fields in this model.

Simulations were performed for a selection of sequence lengths *L* and ensemble repetition numbers *R* and only the 40% assemblies with largest peak firing rates were considered as place-field like for further analysis. A first glance at spatial rate maps and directional tuning of example assemblies reveals prototypical place field firing around feature locations and a wide range of head direction sensitivities (Figure 5A, top). Place field statistics was therefore explored more systematically.

Not surprisingly, mean peak rates increase with repetition number *R* and decrease with sequence length *L* (Figure 5B, top), because activity at late sequence states is broadly distributed in the maze. However, and less obvious, the spatial extent of the place fields decreases with increasing *R* as evidenced by an increase of spatial information and a decrease of the sparseness measure with *R* (Figure 5C, D, top). On a second thought, this effect makes sense because assemblies are activated *R* times *prior* to the time step in which they start the sequence and thus their activity is temporally and spatially less bound to the feature node the smaller *R*. In the extreme case, *R* = 1, ensembles late in the sequence are activated once on broadly distributed locations, which leads to a widening of the firing maps. This effect also explains that place fields with *R* = 1 show much less directional tuning then those with higher *R* (Figure 5E, top). Directional information close to the feature node is carried to distant locations by larger repetition numbers *R* and thus induce directionality.

The fraction *>* 0.5 of direction-tuned assemblies, though it certainly reflects the designed directionality of the features, is relatively high as compared to numbers reported from rat hippocampal area CA3 (14%) and gerbil CA3 (46%) using the same method (Mankin et al. 2019), (for CA1, see (Acharya et al. 2016)). To explain this deviation, I next explored the possibility that additional learning of distinct sensory inputs from non-feature nodes could lead to decreased fractions of directional assemblies. To this end, so-called sensory activations (see Methods Open Field) were added for *Q* = 2 or *Q* = 4 time steps subsequent to the time step in which a theta sequence was first started by a specific ensemble (Figure 5F). Additional sensory activation did not change basic place field statistics, nor their dependence on *R* and *L* (Figure 5B-D), however, as expected, the fraction of direction-tuned assemblies was reduced. Thus learning of direction invariant representations of non-feature nodes during behavior leads to smaller fractions of direction-tuned ensembles as suggested by experiment. The model thereby distinguishes between strong (salient) sensory features that immediately trigger sequences (and occur sparsely), and weaker (regular) sensory inputs that might modulate sequence activity but are not strong enough to trigger sequences on their own.

## Discussion

Place fields are a major physiological hallmark of the hippocampus and therefore many hypotheses have been made to explain how place-field activity and place cell sequences facilitate navigation (Samsonovich and McNaughton 1997; Redish and Touretzky 1997; Battaglia and Treves 1998; Tsodyks 1999; Pfeiffer and Foster 2013; Ponulak and Hopfield 2013; Wu et al. 2017), or how place fields are formed by integration of multimodal sensory inputs (Gothard et al. 1996; Haas et al. 2019; Gauthier and Tank 2018). Specifically the effect of the placement of landmark cues on place field activity are well investigated in classical models (Touretzky and Redish 1996; Redish and Touretzky 1997; Redish 1999). While in the present model these classical ideas could be well integrated as explanations for how salient landmarks elicit activity at the feature nodes, preplay of future behavior (Dragoi and Tonegawa 2011, 2013) is not playing a role in any of the previous models. Only recently it was proposed in (Liu et al. 2018a) that prewired sequences, which are extended during behavior would provide a highly efficient predictive representation for navigational learning.

The model presented here extends the ideas of (Liu et al. 2018a) in that it uses the concurrently active sequences that are linked to salient sensory features to efficiently encode space, while providing a neurodynamical substrate for path integration that is not explicitly dependent on the integration of velocity vectors. The idea that accurate spatial navigation is not tightly linked to physical movement but rather relies on propagation in an abstract space is supported by classical work from (Mittelstaedt and Glasauer 1991) reporting that humans and rodents are relatively imprecise in path integration during 2-dimensional homing. Also psychological experiments showing that spatial maps are rather topological than metric (Warren et al. 2017) fit into this picture.

A feature of the present theory is that it naturally links spatial representations to action selection via reinforcement learning. Whereas (Foster et al. 2000; Fremaux et al. 2013) already proposed a connection between the hippocampal place code and reinforcement learning without activity sequences, the present paper also provides a functional interpretation for sequence replay as facilitating the consolidation of action memory. This interpretation is in line with theories proposing striatal learning to be involved in goal-directed navigation (Johnson et al. 2007) and explaining predictive striatal reward activity (van der Meer and Redish 2009) as a result of the learning of action parameters *m*_*j*_ induced by hippocampal sequences. A direct comparison to reinforcement-based navigation models without preplay (Foster et al. 2000; Fremaux et al. 2013) is, however, not straight forward, since the elementary units (place cells vs. sequences) are fundamentally different for both approaches and thus scaling up one model can always lead to a better performances. A fair comparison could be achieved in terms of computational time for equal performance. This is an interesting avenue to explore for later studies, particularly when also taking into account different neuromorphic hardware implementations (Friedmann et al. 2013).

Hippocampal replay has been proposed to underly planning of trajectories in a known environment (Pfeiffer and Foster 2013; Wu et al. 2017), since sequences were found to predict future behavioral choices. Also sharp wave ripples, which are associated with sequence replay are decreased when an animal exhibits uncertainty in behavioral choices (Papale et al. 2016). Such findings are generally interpreted to reflect mental travel (Ponulak and Hopfield 2013) before an animal decides on the trajectory to take. The present model offers an alternative interpretation of such awake theta sequences, viz. identification of the feature node that serves as entry to the sequence: Assuming that the combination of landmarks, internal motivation and context sets an excitability bias in the sequence generating network (see Appendix A), the sequences that are about to be started once the animal starts the behavior will be more likely activated offline spontaneously already beforehand. In this interpretation, vicarious trial and error behaviors, in which animals are uncertain about a decision, are associated with a reduced excitability of the system, not allowing sequences to evolve. Similarly, errors in sensory processing, or in the representation of contextual information will lead to triggering of wrong or no sequences and thereby induce a correlation between behavioral success and sequence replay. Most importantly, in the framework of this model the link between decision making and offline replay is correlational and not causal. Behavioral decision making in downstream brain areas and offline sequences are linked by a common cause, which is the activation of the correct sequence. The increased excitability of perpetually activated sequences would also explain their abundance in rest periods after strongly repetitive behaviors (Jackson et al. 2006; O’Neill et al. 2006; Singer and Frank 2009).

A potential criticism to the model arises from physiological reports that offline sequences are plastic and change from preplay to replay mostly by adding new neurons (Grosmark and Buzsaki 2016; Liu et al. 2018a; Chenani et al. 2019), whereas in the present model the sequences remain fixed. There are two ways to explain sequence plasticity in the realm of this model. First, sequences plasticity could be an epiphenomenon resulting from replay of multiple sequences. Instead of replaying sequences individually in the serial order of their appearance (as used for the simulations in this paper), multiple sequences can also be replayed in parallel with the same temporal relation as during the search task. Such parallel replay would increase the number of active neurons as compared to preplay and even further improves model performance (Appendix C), making it the preferred explanation of neuronal plasticity during replay. Second, as proposed by (Liu et al. 2018a) sequences may only be prewired with short lengths and the long sequences required in the present model would then be generated on the fly by flexibly connecting short sequence snippets. Also this idea explains sequence plasticity since offline replay would activate the newly generated long sequences, however, it is unclear what could be the mechanisms underlying the on-demand concatenation of snippets. The latter idea cannot be directly modelled by the present theory since it does not provide explanations of how or when long sequences are be prewired.

The three simulated scenarios (random maze, alternation task, open field) presented in this paper arguably only represent a small subset of studied navigational behaviors. Further model extension are therefore necessary to fully evaluate this sequence-based approach to the hippocampal space code. A natural next step would be to more specifically study the effect of less artificial geometric placement of visual features in open fields on place field activity as it was done in classical non-sequence based models (Touretzky and Redish 1996; Redish and Touretzky 1997). Also adding motivation for explaining free exploration strategies (Eilam and Golani 1989; Benjamini et al. 2011) without physical rewards could be implemented by adding intrinsic reward terms based on novelty.

First steps towards a biological implementation of network mechanisms accounting for how sensory inputs can be linked to prewired sequences by synaptic learning have been sketched in Appendix A. The main problem that such models need to solve is how the sequences in each theta cycle are moved forward in register with the inputs, i.e., what is the mechanisms the let the starting ensemble of the theta sequence progress in alignment with behavior. The model explored in Appendix A, which achieves this progression by an STDP rule with a depression at small positive lags, already exhibits some interesting features mentioned in experimental reports. First, the place fields tend to get larger further into the linear track (Gothard et al. 1996; Haas et al. 2019), indicating that fields late in a sequence accumulate more sensory information. Second, Liu et al. (2018a) found evidence for prewired sequence motifs being shorter than the observed sequences. The length of the sequences in the model of Appendix A varies with mean synaptic weights and as a function of all other parameters defining neuronal excitability, hence, arguing that transitions from short to long sequences can result from general excitability increases.

The model presented in this paper makes two specific experimental prediction by which it can be falsified. 1) Sequences should have direction-dependent components. According to Figure 5F sensory spikes occurring early in the theta cycle would be direction-insensitive, whereas the late spikes that represent the prospective sequences should have a preferred head direction. This prediction is in line with findings for hippocampal area CA1, where phase precession was shown to be direction dependent (Huxter et al. 2008), however, so far there are no results on the directional dependence of sequential correlations in certain theta phases, yet. Similarly, rats with dentate gyrus lesions were shown to lack prospective firing in area CA3 (Sasaki et al. 2018), showing that place field activity may have different input contributions depending on theta phase, consistent with earlier CA1 models (Chance 2012) and findings suggesting theta phase specific inputs to CA1 (Colgin et al. 2009; Schmidt et al. 2019). 2) The model explicitly assumes that replays are elicited by reward. It is well known that rewards strongly correlate with the incidence of replay events (Singer and Frank 2009; Fernandez-Ruiz et al. 2019), however, it is unclear whether replays that still occur at unrewarded locations are due to a specific reward expectation (as would be predicted by the model) or can also be induced by reward-unrelated factors.

There is one additional prediction that is not strictly required for the model to hold, but would strongly corroborate the main idea that sequences are triggered by salient features. To efficiently use the dynamical reservoir of the sequence generating network, long sequences are only required in sparsely ornamented mazes (Figure 3F). In spaces with many features, sequences need to bridge less distances and thus finding that sequence lengths are indeed reflecting the sensory richness of an environment would strongly argue in favor of a dynamic process regulating sequence lengths in a task-dependent fashion. There is already some experimental evidence fitting this prediction for very long sequences from running wheels and treadmills (Pastalkova et al. 2008; Malvache et al. 2016), which should represent the most feature-less behavioral situations. Also findings that sequence lengths within a theta cycle correlated with movement velocity (Gupta et al. 2012) and distance to goal (Wikenheiser and Redish 2015) corroborate the existence of dynamical sequence length adaptation. A specific test, however would require recording sequences in parallel to a controlled reduction of the number of features.

Cortical networks have been described to show stereotyped spontaneous activity patterns on multiple temporal and spatial scales (Raichle et al. 2001; Kenet et al. 2003). Studies on primary visual cortex have shown that these spontaneous patterns carry over to stimulus processing (Kenet et al. 2003; Smith et al. 2018) and thus the idea that intrinsic prewiring can be useful for neural information processing is not particularly special for the hippocampus, even more so as sequence like activity can develop in recurrent networks spontaneously (Kitano et al. 2002; Aviel et al. 2003). It is less obvious, however, whether there is a common computational principle favoring dynamical prestructure. One promising candidate theory could be “reservoir computing” (Maass et al. 2002; Jaeger and Haas 2004), which has convincingly shown that a rich general purpose dynamics in neural networks can be used to represent arbitrary input-output relations in time (Legenstein et al. 2003; Sussillo and Abbott 2009) by means of learning feed-forward (and sometimes also feedback) synaptic weights. The theory laid out in this paper suggests that hippocampal sequences may represent such a dynamical reservoir and at the same time could serve as a blueprint for how such neuronal reservoirs could be made use of in other cortical areas, which remains for further exploration.

## Acknowledgments

The author is very grateful to George Dragoi for comments on the manuscript. The author declares no conflict of interest.

## Data availability

The manuscript only contains simulated data. All software used to generate the data will be made available upon reasonable request to the author.

## A Biological feasibility

To show that sequence propagation in coordination with movement-induced temporally evolving sensory input can be biologically plausible with spiking neurons, a model of 64 exponential integrate-and-fire-neurons was implemented that excite each other in two sequences. The first sequence from neuron 1 to neuron 53 is triggered by the start at position 0 of 150 cm linear track and the second sequence (54 to 64) is triggered by reward expectation at position 120 cm. Importantly, a sequence, after having been started by landmark input, would terminate prematurely after the first theta cycle if not sustained by other synaptic inputs.

The simulations are summarized in Figure 6 (details are outlined in a Methods part below). In brief, neurons are assumed to be connected such that they excite each other in sequence during one theta cycle. Once a strong landmark input starts the first sequence at the beginning of the track a position-dependent sensory signal is transformed to a second set of synaptic conductances that evolve in time in parallel to the intrinsically wired sequential activation of neurons. By means of a spike-timing dependent learning rule the initially weak sensory inputs get strengthened by the sequence induced postsynaptic spikes such that once the strong landmark-induced input is gone (after the first theta cycles), the learned sensory inputs take over and keep the sequence alive over multiple theta cycles (Figure 6A,B). Averaging over multiple trials then results in spatial firing maps that extend along the linear track (Figure 6C) and that allow to decode the position of the agent. One the theta time scale, the sequential activity structure by construction translates to phase precession with respect to the theta-rhythmic input current in each individual neuron (Figure 6D).

**Figure 6:**
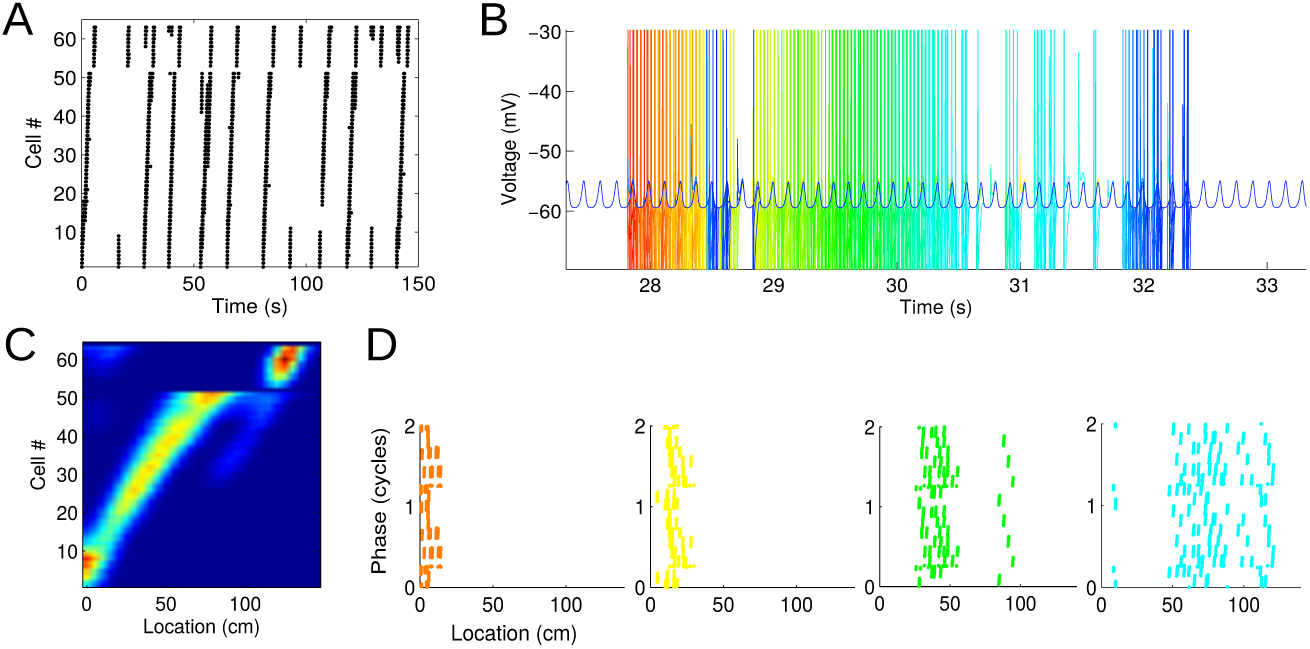
Sequences from simulations of spiking neurons. (A) Spike raster plot for 13 runs of a simulated agent on a linear track. At the beginning of each run an initial sequence of about 7 to 8 neurons is triggered. Close to the reward site a second sequence is triggered. (B) Voltage traces of exponential integrate and fire neurons for the third run as an example. The different colors indicate different neurons. All neurons receive the same inhibitory theta rhythmic input. (C) Firing rate maps (place fields) of all neurons derived from the 13 runs shown in A. (D) Phase precession plots for four example neurons (colors as in B).

Phase precession also indicates that each neuron fires at multiple theta cycles and at every cycle a different neuron leads the theta sequence (see also main Figure 1C). This cycle-by-cycle shifting of the sequence start is only possible if the neuron that starts the sequence in one cycle stops firing in the next cycle thus allowing the sequence to synchronize with the movement induced propagation of sensory inputs. In the model shown in Figure 6 the cycle-by-cycle shifting of the sequence start is implemented by spike-timing dependent plasticity rule that depresses synapses if pre- and postsynaptic activity have small positive lags, such that in the next cycle the sensory synapses transmitting information from the past are not longer strong enough to start the sequence. The feed-forward synaptic matrix thus does not convey a static map of the sensory environment but dynamically changes cycle-by-cycle thereby encoding the position of the agent.

### A.1 Simulation details

The subthreshold membrane potential *u* (in mV) of each of these neurons evolves according to

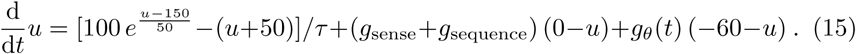

yielding a resting potential a little lower than −50.

If the voltage reaches its individual cut-off potential *u*_*c*_ a spike is triggered and the voltage is reset to − 0.2. After each spike the neuron is refractory for 2 ms. The integration time constant is taken as *τ* = 50 ms. Note that the model is formulated such that the synaptic input conductances have units 1*/*time (e.g., kHz).

The cut-off potentials *u*_*c*_ are randomly drawn from the uniform distribution in the interval [−50, −30].

The conductance *g*_*θ*_(*t*) imposes an inhibitory theta rhythm with amplitude 0.4 kHz and a running speed (*v*) dependent frequency

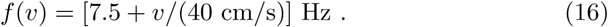

The excitatory synaptic conductance is a sum of sensory inputs *g*_sense_ and sequence generating inputs *g*_sequence_.

The sensory inputs are modeled as a weighted sum of individual stimulus-induced components

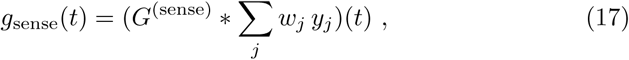

that reflect the sensory inputs on a simulated 1.5 m linear track. The convolution kernel

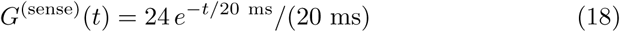

implements synaptic filtering, the weights *w*_*j*_ are resulting from synaptic learning (see below), and *y*_*j*_ represent sensory input channels that are the projections on the principal components of 1000-dimensional sparse sensory activity vectors 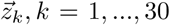. The number of principal components is restricted to capture 85% of the variance.

The sparse sensory activity vectors are generated as

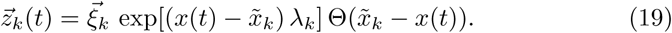

Here, *ξ*_*k*_ is a 1000-dimensional binary random vector with 1% of the entries set to 1. The positions 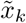 of the sensory stimuli are equidistantly placed on the track, whereas the decay rates *λ*_*k*_ of the stimuli are randomly drawn from the interval between 5 and 35 cm. For visual stimuli, the decay constants *λ*_*k*_ can be interpreted as how visible (or how large) they are. Θ denotes the Heaviside step function.

The sensory model is driven by the running trajectory *x*(*t*) of the simulated agent on a one-dimensional linear track. Random trajectories are derived from the acceleration profile

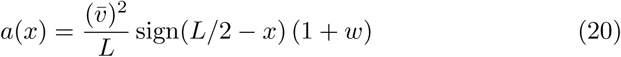

where *w* is a random variable distributed according to 𝒩 (0, 0.3) and refreshed every simulated 200 ms, *L* = 150 cm is the track length, 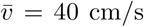 is the mean speed, and *x* is the position on the track.

The sequence conductances *g*_sequence_ are derived from

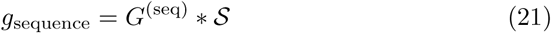

with the synaptic filter kernel

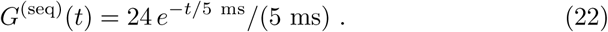

The sequence input S is set to 1 for 2 ms in neuron *n* after neuron *n* − 1 elicited a spike. Sequences are started at neuron 1 in the beginning of a trail by a 2 ms “landmark-induced” conductance of 120 kHz. A second “landmark-induced” conductance is delivered to neuron 53 when the simulated animal has reached the last 20% of the linear track indicating reward expectation.

#### Synaptic Plasticity

The synaptic weights were updated while the agent runs on the linear track according to the following spike-timing dependent learning rule.

the input *y*_*j*_(*t*) is linearly filtered by the kernel

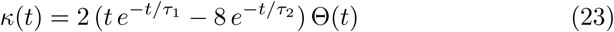

for each principal component *j* independently (time constants were *τ*_1_ = 30 ms and *τ*_2_ = 100 ms). The resulting presynaptic traces *π*_*j*_ = *κ* * *y*_*j*_ are then used to compute weight updates Δ*w*_*j*_ at the time *t*_*p*_ of a postsynaptic spike by

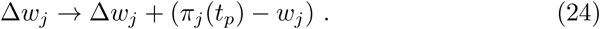

All 100 ms the weight updates Δ*w*_*j*_ are added to the respective weights *w*_*j*_ and set to zero afterwards. After every weight change, weights are clipped at an upper bound of 1 and a lower bound of 0. The learning procedure thus implements a variant of a time-averaged spike-timing dependent plasticity rule.

## B Decay of eligibility trace

Here the effect of the forgetting constant *γ* of the synaptic eligibility trace was explored for navigational learning either without or with replay (Figure 7 Left). Generally, the relative duration after 150 trials does not depend much on *γ* in the case with replay achieving good performance even for relatively fast forgetting (*γ* = 0.25), whereas without replay attenuation improves below 0.4 only for very slow forgetting constants of larger than 0.95. The small differences in performance due to different *γ* values are particularly remarkable since the sparseness (Treves and Rolls 1992) of the weight prediction vector *w* massively depends on *γ*, while sparseness scores increase only little with replay (Figure 7 Right). As expected, slow forgetting (large *γ*) leads to densely populated vectors (high sparseness score) and fast forgetting (small *γ*) leads to sparse vectors (low sparseness score). This finding suggests that relative duration only little depends on the structure of the reward prediction weights and is mostly improved by learning of action parameters during replay.

**Figure 7:**
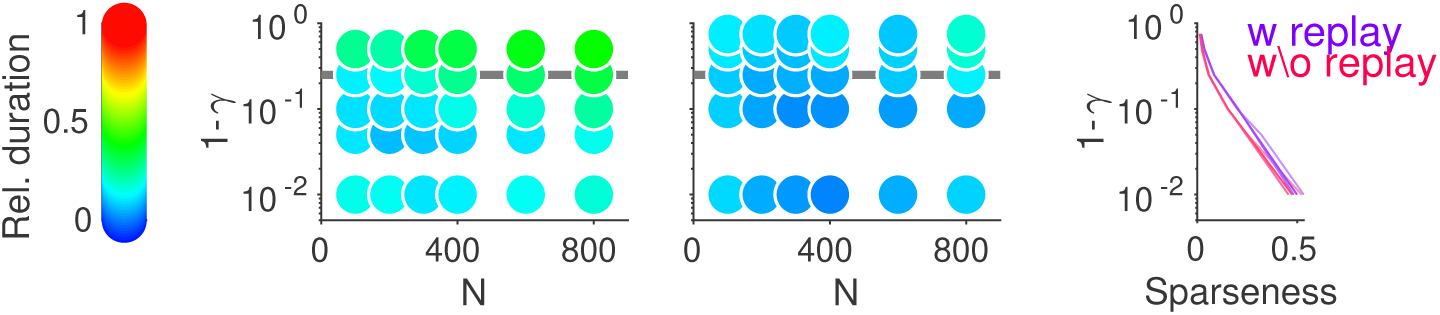
Left: Relative duration (color code as indicated) as a function of forgetting constants *γ* and maze size without replay session (left) and with replay sessions (right) after 150 trials (cf. Figure 3C cyan). Horizontal grey line indicates *γ* = 0.75, the value used in all simulations except here. Right: Sparseness of the reward prediction weight vector (*w*_*n*_) as a function of *γ* (purple and red lines from simulations with or without replay as indicated).

Thus replay allows for a substantial shortening of memory time scales of the eligibility trace.

## C Replay of Multiple Sequences

A potential concern about the validity of the presented model, requiring many long sequences to be concurrently expressed in the network, could arise from capacity considerations, i.e., the number of different sequences that can be played out at the same time could just be too small to map navigational problems of real-life complexity. However, capacity estimates outlined in Appendix D show that a hippocampal CA3 area with about 500,000 neurons (bilaterally) could easily host few thousands of distinct sequences. Moreover, slow homeostatic reorganization of the recurrent sequence-producing circuits could maintain the network in a state that always comprises sufficiently many new sequences at the cost of loosing old ones (Medina and Leibold 2014). Such a palimpsest property would require that navigational strategies in very familiar environments should be represented in different brain structures and therefore potentially also follow different rules than those for novel environments.

In the standard version of the model sequences are replayed serially after each trial in the same order as they have been activated during the search. However, physiological evidence suggests that sequences are plastic in the sense that sequences observed at reward sites have additional neurons included as compared to preplay sequences (Grosmark and Buzsaki 2016; Liu et al. 2018a; Chenani et al. 2019).

In the present model this plasticity can be interpreted in two ways (see Discussion). One of these interpretations suggests that prewired sequences are not just replayed in isolation, but sequences are replayed in parallel with the same temporal relation as they occurred during the search task leading to increased participation of neurons during replay episodes.

To test, whether such parallel replay would still work in the model, I compared mean search durations between the two types of replay in an example parameter regime (*L* = 150, *R* = 7, *N* = 400) and found that while the asymptotic (late) relative durations do not significantly differ, performance for parallel replay seems to converge faster, since relative durations for early trials (11 to 60) are significantly shorter for parallel replay and not significantly different to late trials; Figure 8.

**Figure 8:**
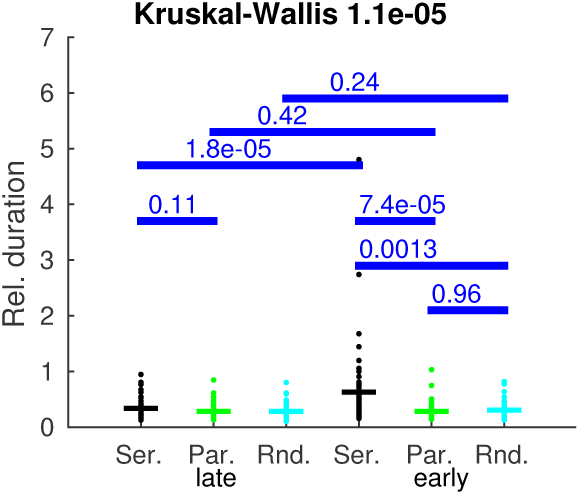
Parallel replay of sequences converges faster than serial replay. Shown are mean relative trial durations for 50 random maze search tasks comparing serial (Ser.: black) parallel (Par.: green) and randomized parallel (Rand.: cyan) replay for early trials (right: 11 to 60) and late trials (left: 101 to 150). All p-values (blue) are derived from two sided signed rank tests. Bars indicate the means.

In a further set of control simulations, the starting sequence was selected randomly from the set of experienced sequences and after that sequences were again replayed in parallel with the same temporal relation as they occurred during the search task. Also for this randomized protocol duration were the same as for the non-randomized parallel replay protocol.

## D Capacity for Sequences

The sequence length *L* is a main parameter determining, how well the model learns a navigational task. Moreover, a sequence is triggered whenever the agent arrives at a feature node and thus several sequences will be active at any point in time. A necessary requirement for the model to be biologically plausible is thus that the putative hippocampal CA3 network that hosts the many sequences can play them out reliably whenever demanded. In this section I provide an assessment of how many sequences can be stored in a network, showing that the capacity for sequences is very large and biology thus presumably does not impose to strong constraints on the sequence generating network. The theory outlined below follows previous work in (Willshaw et al. 1969; Leibold and Kempter 2006).

Let us assume *U* neurons, each of which can be synaptically connected to *c*_*m*_ *U* others. The potential connections can be active (synapse exists) or inactive. Only active connections can transmit signals and the synaptic connectivity pattern underlies the stored sequences.

Sequences consist of an ordered activation of neuronal assemblies. A neuron can participate in multiple assemblies independently. The probability by which a neuron is assigned to any given assembly is *ϕ*, such that *M* = *ϕ U* neurons are active in every assembly (A generalization of the theory to assemblies with varying *M* van be found in (Medina and Leibold 2013)). If *S* sequences of length *L* + *R* are stored in the network, then *P* = *S* (*L* + *R* − 1) is the total number of transitions between assemblies the network must be able to realize. Following (Willshaw et al. 1969), the probability *c* of an active synapse is determined by *P* and *ϕ* according to

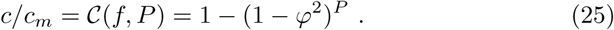

That means that for large *P, c* converges to *c*_*m*_ and all synapses are active. However, if too many synapses are active also random pattern elicit sufficiently strong depolarization and the network will display ongoing random activity. In order to find a maximal *c* for which sequences can still be properly replayed, I use an approach motivated by signal detection theory as in (Leibold and Kempter 2006). If *q* sequences are concurrently active in the network, the noise level (average input upon a random pattern of *M* active neurons) will be *c M q*, whereas the signal level (input if only the correct pattern of *M* neurons is active) is *c*_*m*_*M*. Assuming binomial statistics for large *U* and *M*, and introducing a separation parameter *d* between noise and signal distribution, one obtains the following threshold condition

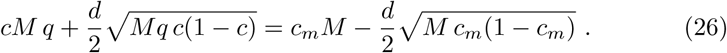

Introducing the abbreviation *x*^2^ = *q c/c*_*m*_ = *q* 𝒞, and considering that the total connectivity needs to be small 0 *< c* « 1, equation 26 is solved by

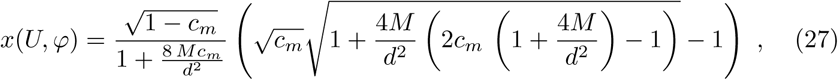

which yields the maximum number of transition between assemblies

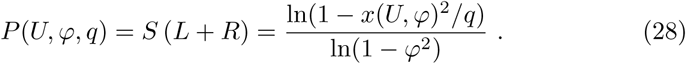

Equation (28) is illustrated in Figure 9 for some parameter examples and *U* = 500, 000 neurons (which approximately corresponds to the bilateral number of CA3 units in rat).

**Figure 9:**
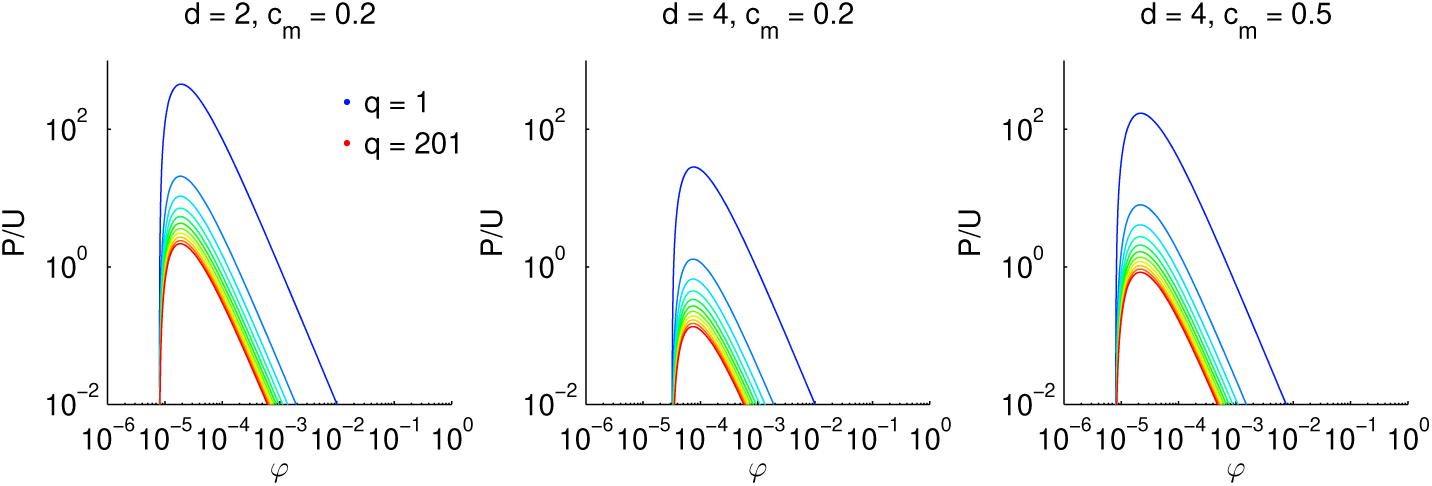
Storage capacity for sequences. Each panel shows the number *P* of associations between assemblies that can be stored in a network divided by the number *U* of neurons for an example parameter regime as indicated on top. The number of neurons is fixed *U* = 500, 000. The ordinate samples coding ratios *ϕ*, which are the fractions of active neurons in an assembly divided by *U*. Colors indicate different numbers *q* of concurrent sequences.

Generally, the decrease in *P* as a function of the number *q* of concurrent sequences is strongest for low *q*. Therefore, many concurrently active sequences still allow for *P/U* ≈ 1 at optimal sparseness levels *ϕ*. A further critical parameter is the signal-to-noise parameter *d*, which should be high to avoid false alarm activation of neurons to ensure persistence of the sequence for many time steps. But even for high signal-to-noise of *d* = 4, assuming a large connectivity parameter *c*_*m*_ = 0.5 can still ensure *P/U* ≈ 1. Thus for *U* = 500, 000, a sequence length of *L* + *R* ≈ 100, the number *S* of concurrently active sequences can reach a value, as high as

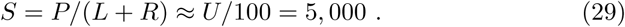

The hippocampus could thus hold few thousand sequences each of which can be started by a salient input feature.

## E Random Mazes

Random mazes are symmetrically connected graphs consisting of *N* nodes. The nodes can only be connected if they are next neighbors on a discrete 2-dimensional square grid.

Mazes are generated recursively. For each newly generated node, each not yet connected neighbor position is connected to the new node with probability *p*. If such a new edge reaches to a neighboring position with an already existing node the edge is made symmetric, otherwise a new node is generated at this position and the same algorithm is applied to the new node. The algorithm starts with a node at coordinates (0, 0) and ends when *N* nodes have been generated. If the recursion terminates before *N* nodes are reached, the recursion is restarted at the node with maximal *x* coordinate.

For small or high values of *p* the mazes generated by this algorithm have elongated and mostly linear structure. For intermediate *p* the graphs contain multiple connected regions that are almost 2-dimensional. All simulations in this paper have been performed for mazes with *p* = 0.5.

## F Movies

### Movie 1: Search of a naive agent

Early example trial of a naive agent navigating an example random maze with a single reward placed at the position of the yellow triangle. The moving black dot indicates position of the agent. Black ticks indicate walls that the agent cannot pass. The colored circles in the center of each node indicates the reward prediction weight of the node (minimal to maximal: blue to red). Using the same color code, the ticks connecting the nodes represent the action parameters *m*_*j*_ corresponding to the four possible movements.

### Movie 2: Search of a trained agent

Late example trial of the same agent as in movie 1.

### Movie 3: Replay in a naive agent

Early replay session of the agent in movie 1.

### Movie 4: Replay in a trained agent

Late replay session of the agent in movie 1.

### Movie 5: Learning of Figure-8-Maze

Learning to alternate between two goal locations over 100 trials. The maze is shown in two copies (see Figure 4 in the main text), which represent motivational states *µ* = 1 and *µ* = 2. The moving dot switches between the mazes whenever there is a switch in he motivational state. The learned alternation behavior becomes visible in the last ≈ 10% of trials.

