## Supplementary figures and images for "A Model for Navigation in Unknown Environments Based on a Reservoir of Hippocampal Sequences"

### Movie S1

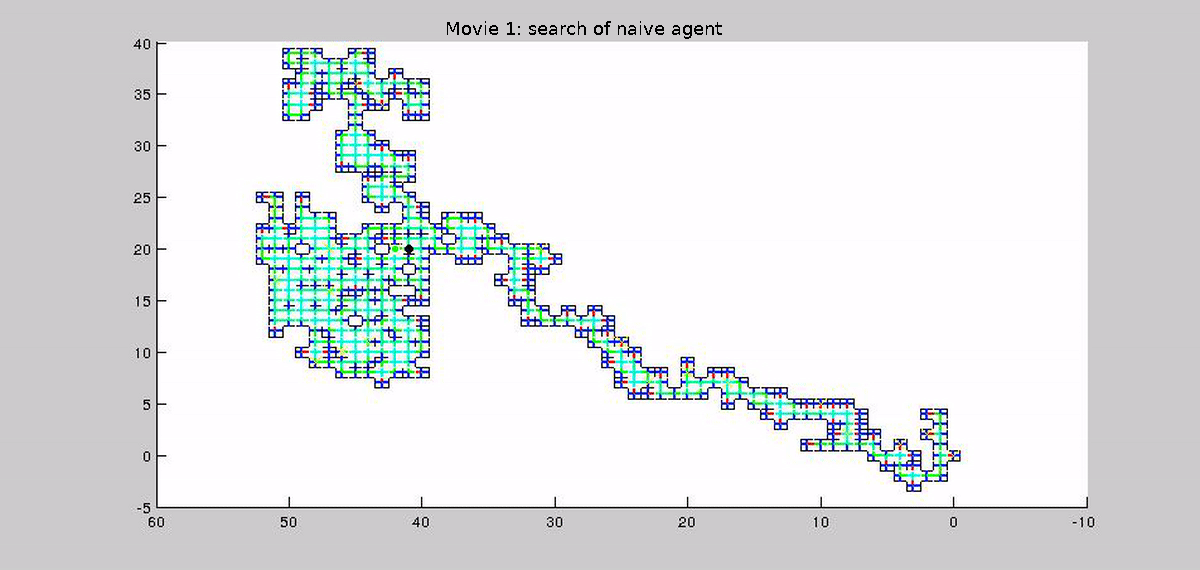

### Movie S2

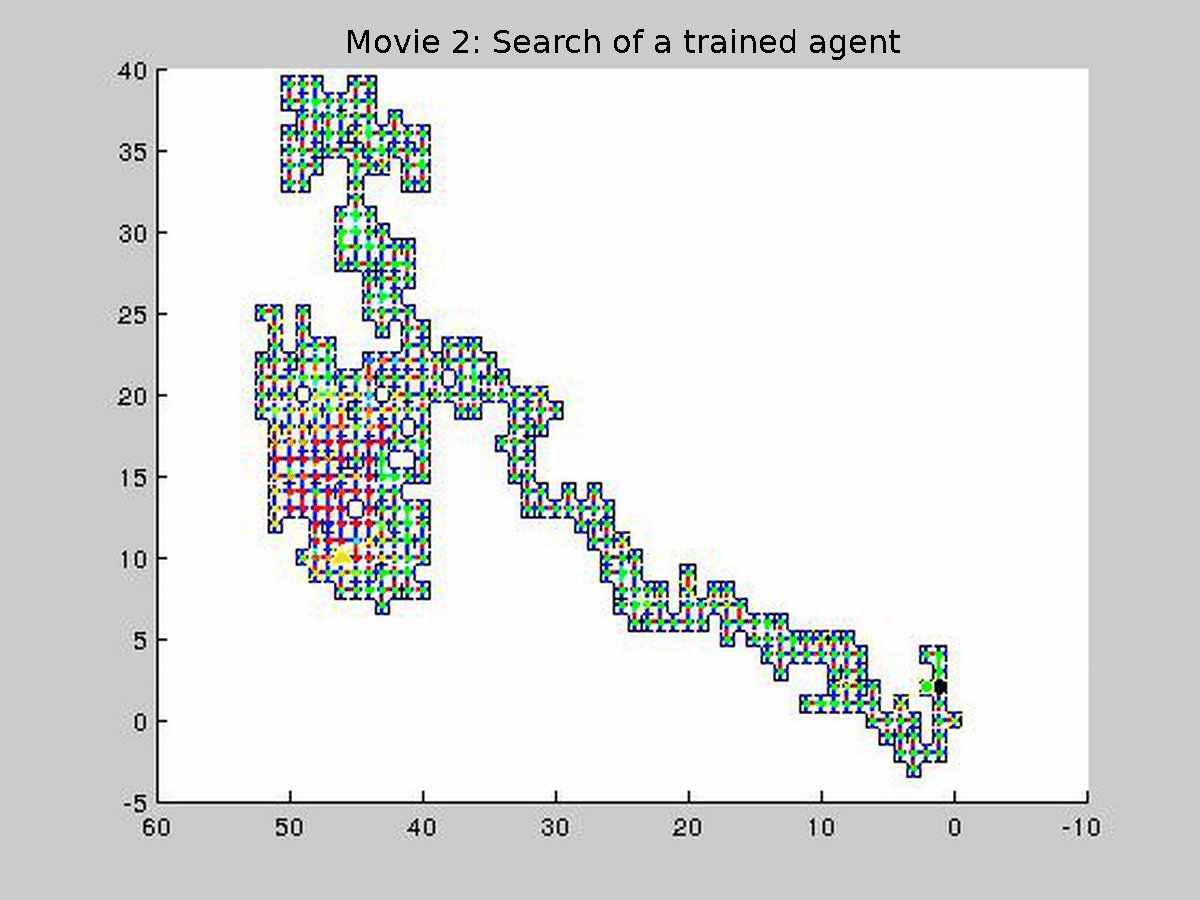

### Movie S3

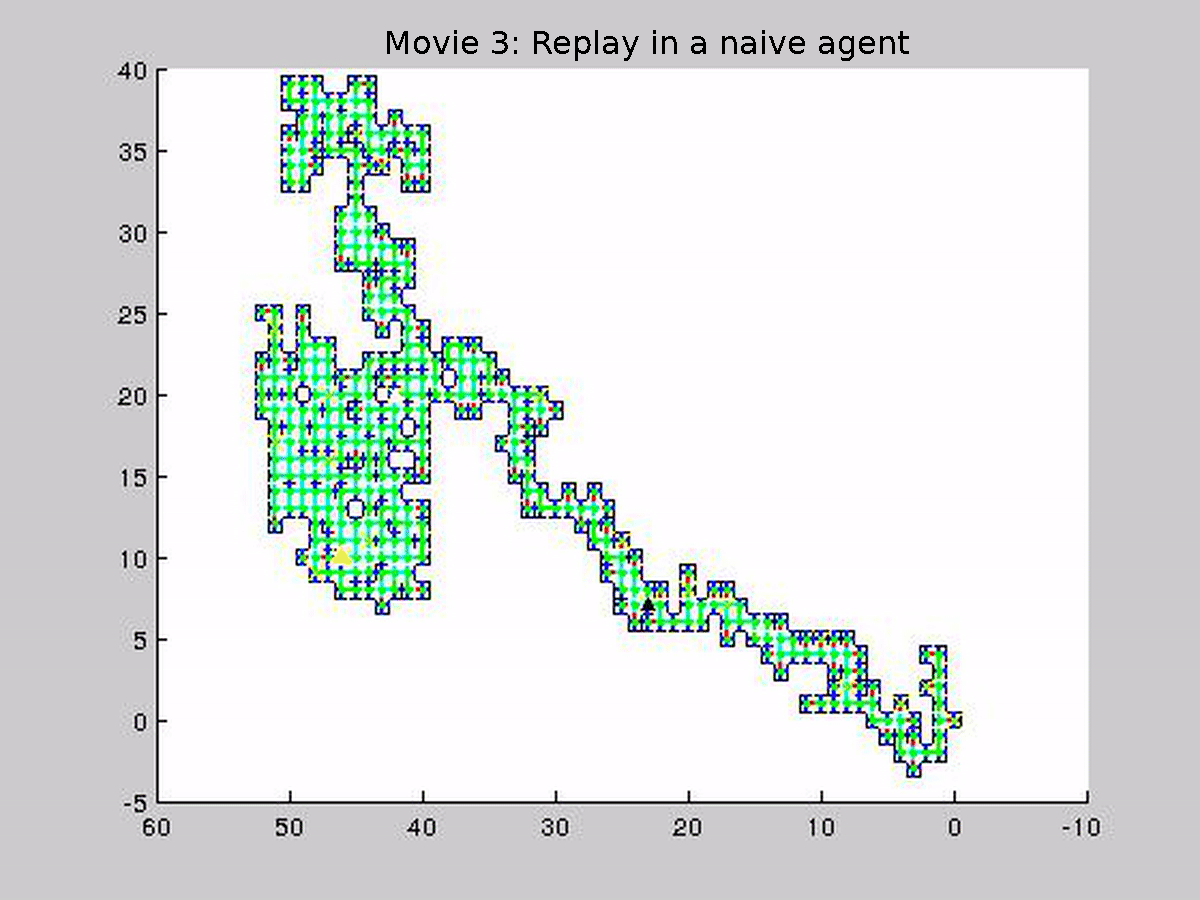

### Movie S4

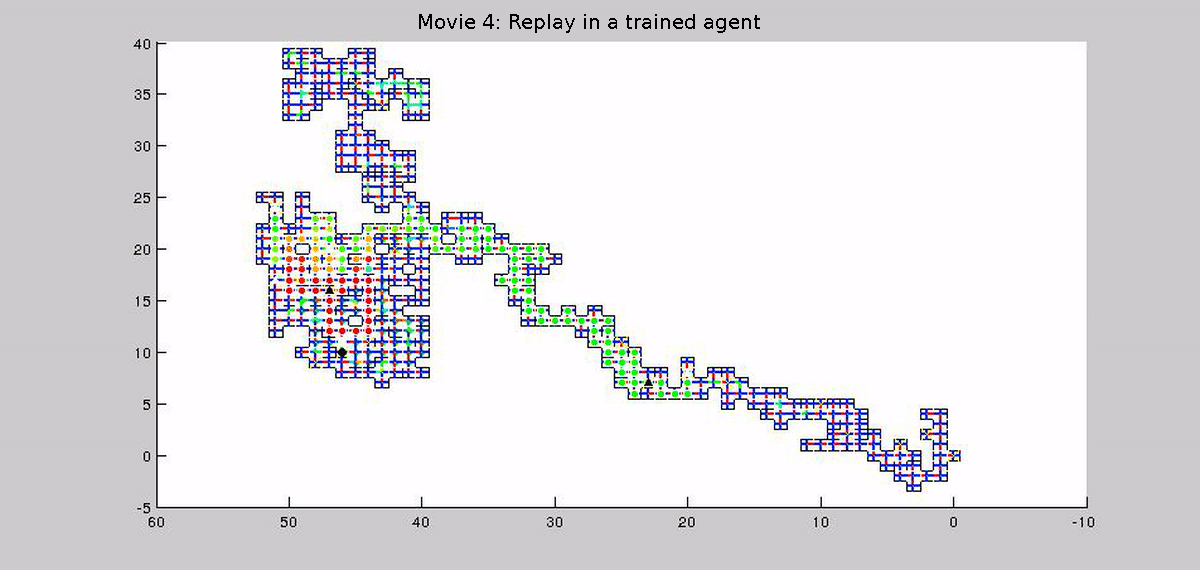

### Movie S5

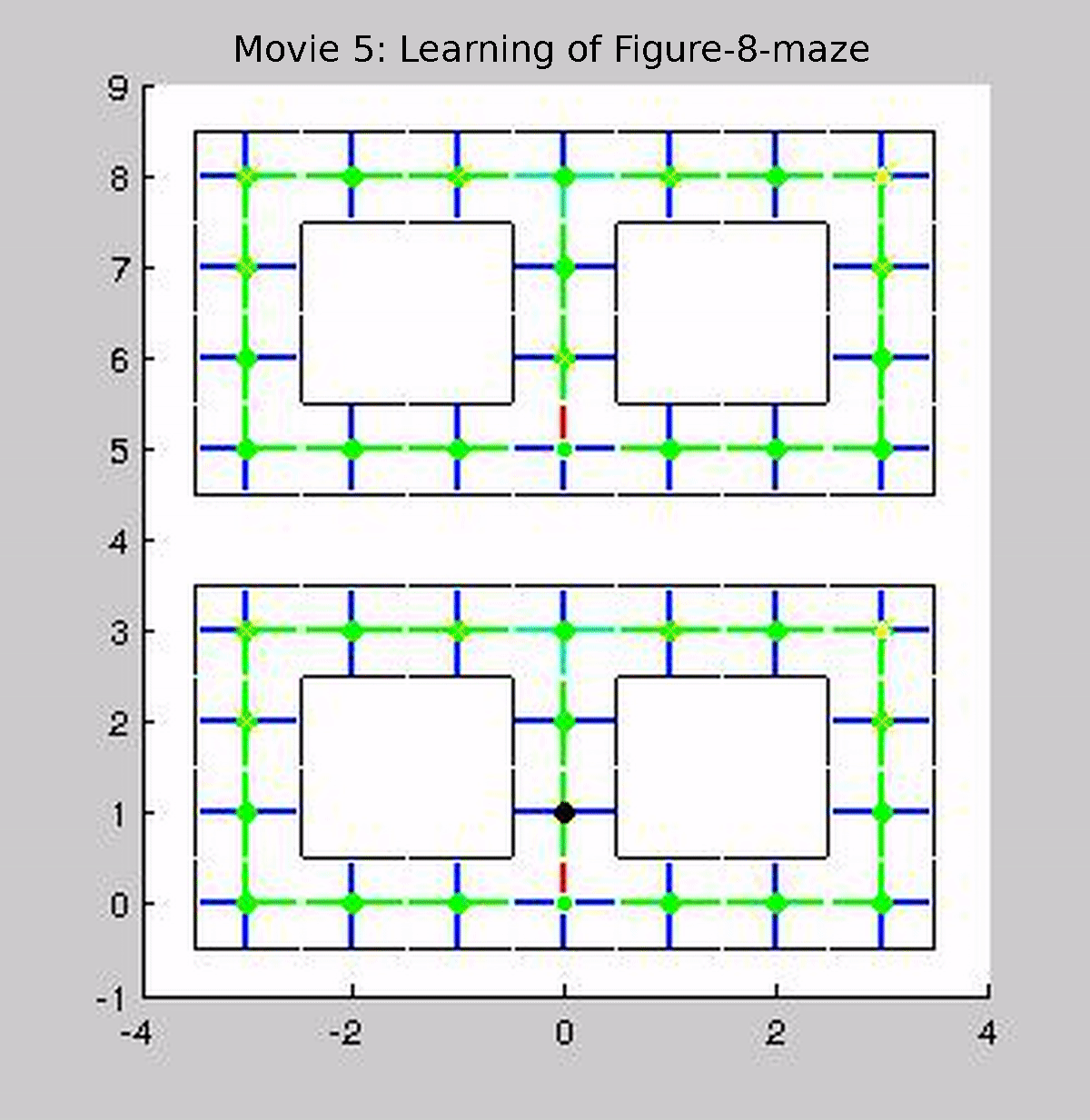
